# XIAP-mediated targeting of endolysosomes to stressed mitochondria occurs in a switch-like, global manner and results in autophagy-independent, sub-organelle level mitochondrial degradation

**DOI:** 10.1101/2023.04.23.538008

**Authors:** Tim Sen Wang, Isabelle Coppens, Nathan Ryan Brady, Anne Hamacher-Brady

## Abstract

Damaged mitochondria can be subject to lysosomal degradation via mitophagy. However, whole-organelle degradation exhibits relatively slow kinetics and thus its impact may be limited in response to acute, fast-acting cellular stress. We previously reported that in Parkin-deficient cells endolysosomes directly target mitochondria when subjected to bioenergetic stress. Here, using high-resolution *live* cell imaging we reveal a striking level of dynamic targeting of Rab5+ early endosomes to stressed mitochondria, culminating in a switch-like accumulation in the entire mitochondrial population, independently of canonical autophagy. This process of rapid, largescale Rab5+ vesicle trafficking to mitochondria coincides with, and is mediated by, XIAP E3 ligase activated mitochondrial ubiquitylation and results in ultrastructural changes to, and degradation of, intra-mitochondrial components. Mitochondria-targeting vesicles include early endosomal subpopulations marked by Rab5 effector APPL1 and ubiquitin-binding endocytic adaptors OPTN, TAX1BP1 and Tollip, and Rab7-positive late endosomes/lysosomes. In Parkin expressing cells, XIAP- and Parkin-dependent mitochondrial targeting and resulting processing modes are competitively regulated. Together, our data suggest that XIAP-mediated targeting of endolysosomes to mitochondria functions as a stress-responsive, sub-organelle level mitochondrial processing mode that is distinct from, and competitive to, Parkin-mediated mitophagy.

## Introduction

Mitochondria are semi-autonomous organelles that form a dynamic, interconnected network through membrane fusion and fission events ^1^. Mitochondria maintain bioenergetic, ionic and metabolic homeostasis, and participate in regulated cell death, inflammatory and innate immune signaling ^2,3^. Consequently, mitochondrial function is vital for organismal health, and mitochondrial dysfunction is linked to numerous diseases, including neurodegeneration and cancer.

It is increasingly appreciated that connectivity among different organelle types is a key component of homeostatic and stress signaling ^4-7^. Mitochondria engage in inter-organelle interactions with multiple organelle types ^8,9^. Membrane contact-based interactions between mitochondria and the endoplasmic reticulum (ER) enable lipid, protein, and ion transfer, and regulate mitochondrial bioenergetics and apoptosis signaling ^10,11^. Furthermore, mitochondria are connected with vesicles of endosomal and lysosomal organelle compartments, referred to here as endolysosomes, through the quality control processes of mitophagy ^12,13^ and mitochondrial-derived vesicles (MDVs) ^14,15^. During mitophagy, dysfunctional mitochondria are sequestered into autophagosomes which sequentially fuse with acid hydrolase-active endosomes and lysosomes for autophagolysosomal degradation. MDVs, on the other hand, form through budding of select mitochondrial contents from the OMM into the cytoplasm and subsequently fuse with lysosomes.

Recent work of ourselves and others has revealed that endolysosomes also directly interact with mitochondria ^16^. Membrane contacts between mitochondria and the lysosome-like vacuole in yeast are under metabolic control ^17^ and serve in phospholipid transport ^18^. In mammalian cells, early endosomes engage in transient “kiss-and-run” interactions with OMMs to transfer iron ^19^, late endosomes/lysosomes regulate mitochondrial morphology ^20^ and related inter-mitochondrial contact formation ^21^, and lysosomes undergo functionally uncharacterized transient interactions with mitochondria ^8,22^. In response to diverse stress stimuli, including BH3-only protein expression and mitochondrial uncoupling ^23^, oxidative stress through laser irradiation or hydrogen peroxide (H_2_O_2_) treatment ^24^, pharmacological induction of intrinsic apoptosis signaling ^25^, and mtDNA damage ^26^, mitochondria are directly targeted by endosomes and endocytic regulators. Moreover, blocking autophagy shifts specific mitophagy programs mediated by Bnip3, Bnip3L/Nix ^23^ and Parkin ^27^ from autophagy to direct endolysosomal targeting modes. These findings suggest a broad and varied interconnectivity between different types of endolysosomes and mitochondria that is at present mechanistically and functionally insufficiently understood.

Here, using high-resolution *live* cell imaging, we investigated the spatio-temporal dynamics and mechanisms regulating endolysosomal interactions with mitochondria, in response to acute mitochondrial bioenergetic and oxidative stress conditions. We reveal extensive and dynamic interactions of endolysosomes with stressed mitochondria and provide mechanistic insights into regulation through X-linked inhibitor of apoptosis (XIAP) E3 ubiquitin ligase and ubiquitin-regulated endocytic adaptors and effectors. The observed rapid, stress-responsive, and autophagy-independent, direct targeting of the entire mitochondrial population by endolysosomes results in a Parkin-alternative mitochondrial processing mode, functioning at the sub-organelle level.

## Results

### Rab5-marked early endosomes accumulate in the mitochondrial compartment in response to various mitochondrial bioenergetic stresses

We previously found that endocytic regulators, including Rab5 and Rabex-5, accumulate in the mitochondrial compartment in response to pro-apoptotic BH3-only protein signaling and bioenergetic stress ^23^ and pharmacological apoptosis induction ^25^. Rab5 targeting of mitochondria was also reported in response to oxidative stress induced by laser irradiation or H_2_O_2_ ^24^. To determine whether targeting of mitochondria by Rab5+ endolysosomal vesicles is a common response to mitochondrial stress, we treated MCF-7 breast cancer cells with a panel of mitochondrial stressors, including nutrient deprivation, ionophores (CCCP, valinomycin, nigericin), and electron transport chain inhibitors (oligomycin and antimycin A), and compared these treatments to H_2_O_2_ and control growth conditions (full medium, FM). Remarkably, all these stressors activated largescale targeting of RFP-Rab5 to mitochondria (**Fig. 1a, b**). Of note, in addition to mitochondrial-accumulated RFP-Rab5, mobile RFP-Rab5+ vesicles were visible in mitochondrial proximity (see color-coded time-sum (Σ(2min)) projections, **Fig. 1a**). In contrast, H_2_O_2_ treatment resulted in mitochondrial RFP-Rab5 accumulation, with a marked depletion of mobile cytosolic RFP-Rab5+ vesicles (**Fig. 1c**). Common to all stressors were significantly increased cellular levels of lipid peroxidation (**Fig. 1d**), and pre-treatment with the antioxidant Trolox abolished valinomycin-induced RFP-Rab5 targeting to mitochondria (**Fig. 1e**). RFP-Rab5 targeting to mitochondria was observed also in HeLa cervical cancer cells in response to valinomycin and nigericin (**Fig. 1f, e**). Valinomycin is commonly applied in the study of Parkin-mediated mitophagy ^28^, unlike CCCP, it does not directly impact lysosomal function ^29,30^, and we here found that it strongly induces mitochondrial targeting by Rab5 in Parkin-deficient MCF-7 and HeLa cancer cell lines. Thus, valinomycin was chosen as the triggering agent for subsequent experiments.

**Fig. 1:**
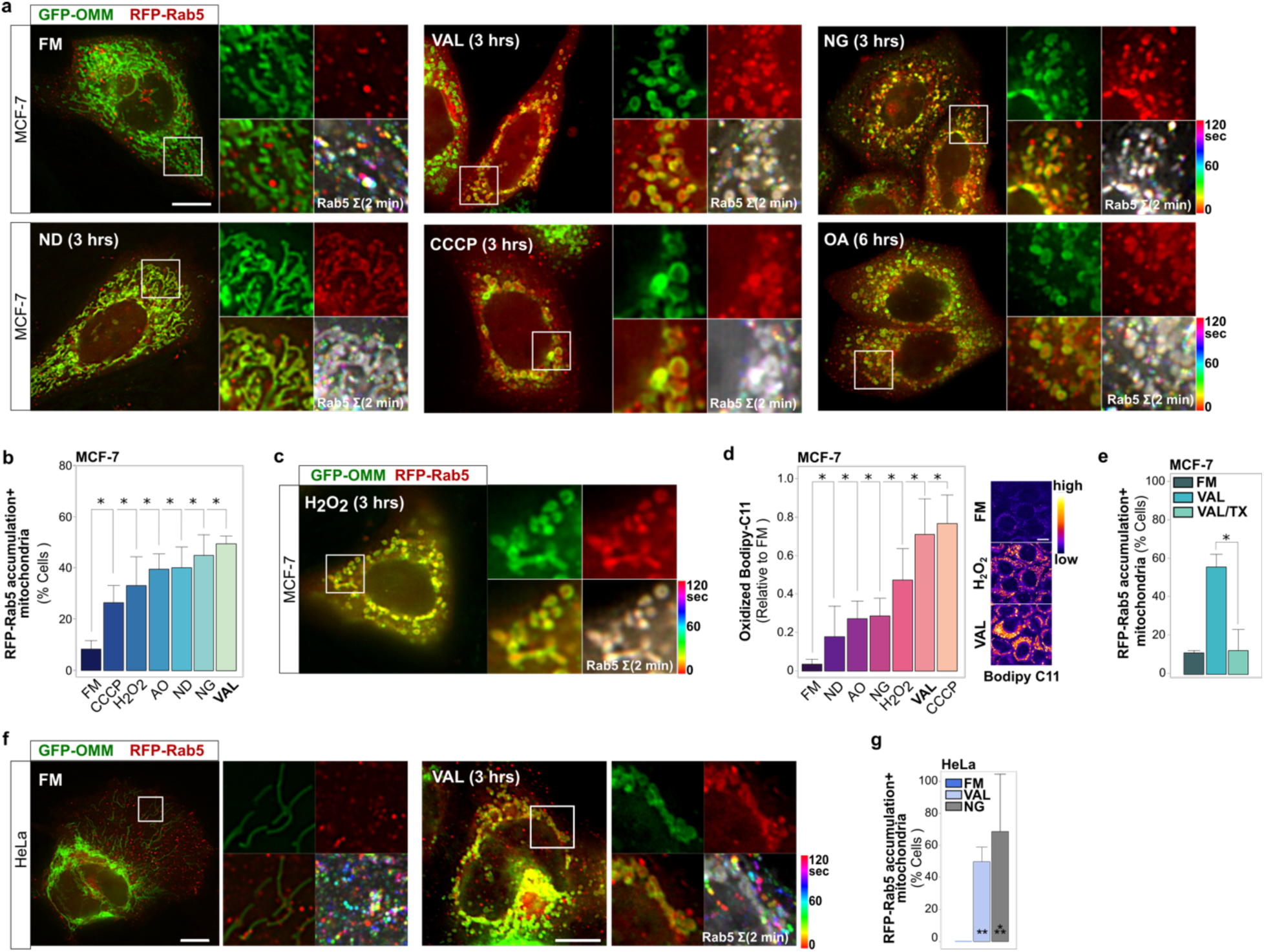
Diverse mitochondrial bioenergetic and oxidative stresses trigger accumulation of early endosomal Rab5 in the mitochondrial compartment. **a** MCF-7 cells co-expressing GFP-OMM and RFP-Rab5 in normal growth medium (full medium, FM), or subjected to nutrient deprivation (ND), valinomycin (VAL; 0.9 µM), CCCP (20 µM), or nigericin (NG; 10 µM) for 3 h, or oligomycin/antimycin A (OA; 10 µM / 25 µM) for 6 h, and then *live* imaged every 5 s for 2 min. Color-merged overview image, single and merged color ROIs, and ROI of 2-min time-projected image of RFP-Rab5+ vesicle movements color-coded by time-step (Σ2 min). **b** MCF-7 cells co-expressing RFP-Rab5 and GFP-OMM subjected to treatments as indicated, scored for mitochondrial compartment targeted by Rab5 (Rab5+ Mitochondria). Graph, mean ± SD of n = 3 experiments. ≥ 200 cells scored per condition. \*\*\**p* ≤ 0.001, \*\*\*\**p* ≤ 0.0001 **c** MCF-7 cell co-expressing GFP-OMM and RFP-Rab5 at 3 h H_2_O_2_ (250 µM). Represented as in (a). **d** MCF-7 cells loaded with Bodipy 581/591 C11 submitted to the indicated conditions. Graph, mean ± SD single cell values of oxidized Bodipy C11 levels; n=3 experiments. Representative images of oxidized Bodipy C11 levels in MCF-7 cells in FM or treated with H_2_O_2_ or VAL displayed in rainbow LUT. **e** MCF-7 cells co-expressing RFP-Rab5 and GFP-OMM treated with VAL (3 h), or pre-treated with 200 µM Trolox (1 h) followed by Trolox/VAL (3 h), scored for Rab5+ mitochondria. Graph, mean ± SD of n = 3 experiments. ≥ 200 cells scored per condition. \**p* ≤ 0.05 **f** HeLa cells co-expressing GFP-OMM and RFP-Rab5 in FM, or subjected to valinomycin (VAL; 0.9 µM) or nigericin (NG; 10 µM) for 3 h, then live imaged every 5 s for 2 min. Color-merged overview image, single and merged color ROIs, and ROI of 2-min time-projected image of RFP-Rab5+ vesicle movements color-coded by time-step (Σ2 min). **g** HeLa cells as in (f), scored for mitochondrial compartment targeted by Rab5 (Rab5+ mitochondria). Graph, mean ± SD of n = 3 experiments. > 65 cells per condition. **p < 0.01, ***p < 0.001 Scale bars, 10 µm.

### Rab5 targeting of depolarized mitochondria occurs in a switch-like and global manner, coinciding with mitochondrial XIAP translocation and ubiquitylation

To capture the spatio-temporal dynamics of largescale mitochondrial Rab5 accumulation, we then performed *live* microscopy of cells that had not yet triggered this phenotype, starting at 3 h of valinomycin treatment (**Fig. 2a, Supplementary Movie 1**). Remarkably, while upon valinomycin treatment the relative temporal onset of accumulation varied from cell-to-cell, RFP-Rab5 accumulated within the entire mitochondrial compartment of all cells in a rapid, switch-like manner. As visible in zoom images (**Fig. 2b**) and intensity traces (**Fig. 2c**), once triggered, the mitochondrial accumulation of RFP-Rab5 occurred over a period of only a few minutes. Following this switch, RFP-Rab5 remained at stably detectable levels within mitochondria.

**Fig. 2:**
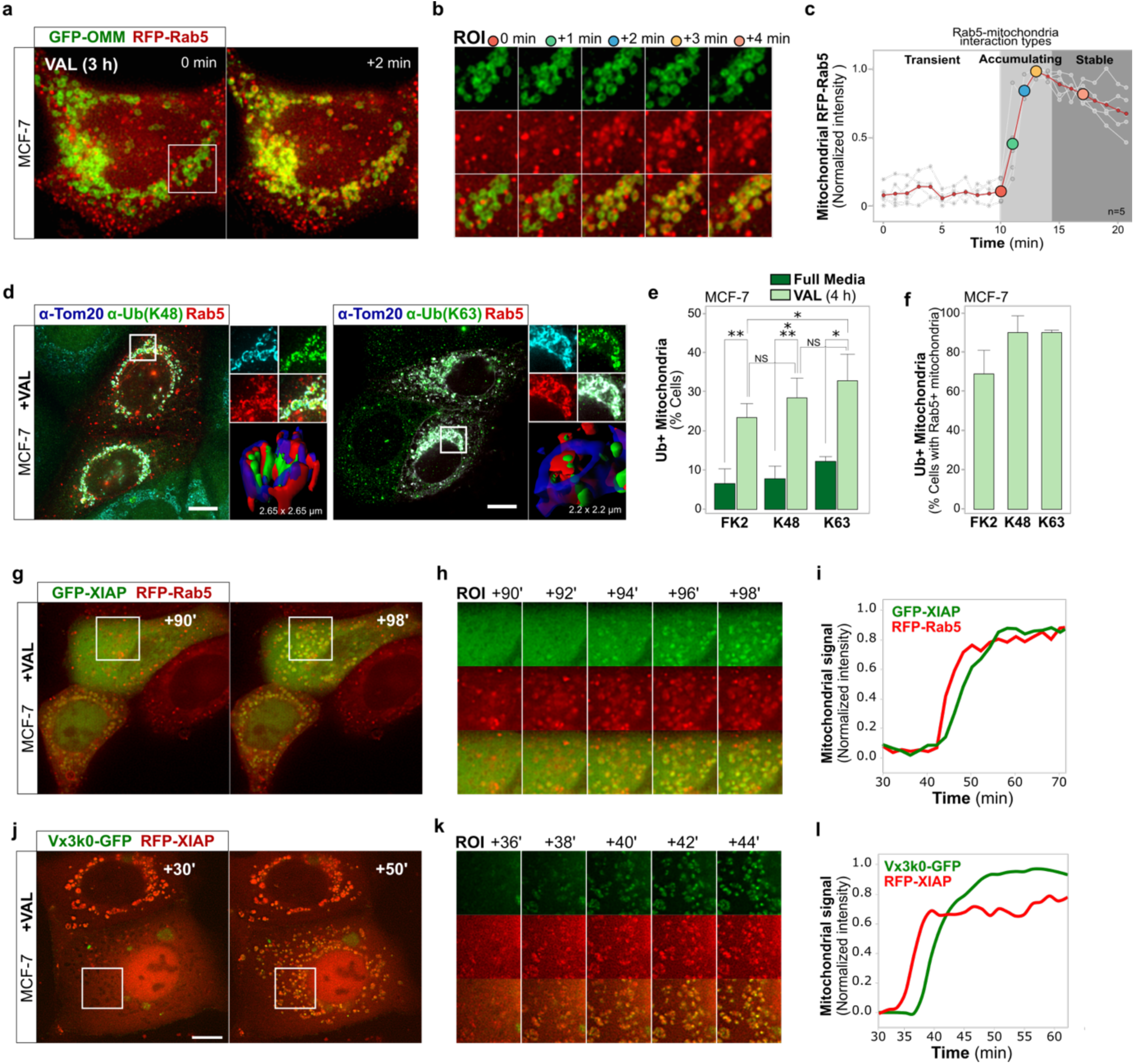
Spatio-temporal dynamics of valinomycin-induced mitochondrial targeting by endosomes, XIAP and ubiquitylation. **a** MCF-7 cell co-expressing GFP-OMM and RFP-Rab5, imaged *live* every 60 s, starting at 3 h VAL. See also **Supplementary Movie 1**. **b** Time-series zoom images of ROI in (a). **c** Quantification of experiment in (a). Normalized intensities of RFP-Rab5 prior to, during, and following accumulation at mitochondria. Overlay of single cell measurements, with start of accumulation phase normalized to timepoint 10 min. Red trace, mean values. Grey traces, individual cells. n = 5 cells analyzed. **d** MCF-7 cells expressing RFP-Rab5, treated with VAL for 4 h. IF of TOM20 and K48- or K63-linked ubiquitin chains. Merged color overview image and ROI zoom images. 3D surface reconstruction of a mitochondrion in ROI. **e** Quantification of experiment in (d) and cells stained for FK2-detectable ubiquitin conjugates. Cells scored for mitochondria positive for indicated ubiquitin IF signals. Graph, mean ¬± SD of n = 3 experiments. > 225 cells per condition. *p < 0.05, **p < 0.01, ***p < 0.005 **f** Quantification of experiment in (d, e). Cells with RFP-Rab5+ mitochondria were scored for mitochondria positive for indicated ubiquitin IF signals. Graph, mean ± SD of n = 3 experiments. *p < 0.05, **p < 0.01, ***p < 0.005 **g** MCF-7 cells co-expressing GFP-XIAP and RFP-Rab5, *live* imaged every 1 min over 40 min, after VAL administration. See also **Supplementary Movie 2**. **h** Time-course zoom images from ROI in (g). **i** Quantification of experiment in (g). Plotted is the appearance of RFP-Rab5 and GFP-XIAP fluorescence at mitochondria. Representative example of n = 5 cells analyzed. **j** MCF-7 cells co-expressing RFP-XIAP and K63-linked polyubiquitin chain sensor Vx3K0-GFP, *live* imaged every 1 min over 50+ min, after VAL administration. See also **Supplementary Movie 3**. **k** Time-course zoom images from ROI in (j). **l** Quantification of experiment in (j). Plotted is the appearance of Vx3K0-GFP and RFP-XIAP fluorescence at mitochondria. Representative example of n = 3 cells analyzed. Scale bars, 10 µm.

We previously reported that the cytosolic E3 ubiquitin ligase and caspase inhibitor XIAP ^31^, under conditions of bioenergetic or apoptotic stresses, localized to ubiquitylated mitochondria marked by endocytic markers ^23,32^. Using ubiquitin chain-specific antibodies, we detected K48- and K63-linked polyubiquitin chains inside mitochondria following valinomycin treatment (**Fig. 2d, e**). Similarly, mitochondrial ubiquitin conjugates were detected by the mono- and polyubiquitin conjugate-specific antibody clone FK2 (**Fig. 2d, e**). Quantification of mitochondrial ubiquitylation within cells positive for mitochondrial accumulated RFP-Rab5 revealed a strong correlation between these two phenotypes (**Fig. 2f**).

We thus performed *live* cell imaging to elucidate the spatio-temporal relationship between XIAP translocation to mitochondria, mitochondrial ubiquitylation, and the above-described rapid endosomal targeting of valinomycin-depolarized mitochondria. Monitoring of GFP-XIAP together with RFP-Rab5 demonstrated that the onset of the switch-like mitochondrial accumulation of RFP-Rab5 coincides with the intra-mitochondrial accumulation of GFP-XIAP, with slightly faster kinetics for RFP-Rab5 accumulation once initiated (**Fig. 2g-i, Supplementary Movie 2**). Simultaneous observation of the dynamics of RFP-XIAP and K63-polyubiquitin chain sensor Vx3K0-GFP ^33^ revealed that XIAP translocation into mitochondria occurred minutes prior to the detection of intra-mitochondrial K63-linked polyubiquitylation (**Fig. 2j-l, Supplementary Movie S3**).

### XIAP E3 ligase activity mediates the largescale mitochondrial targeting by Rab5+ early endosomes in response to valinomycin

Consistent with our prior findings ^23^, K63-linked ubiquitin chains appeared confined to inner mitochondrial compartments. However, FK2 antibody detection of other mono- and poly-ubiquitin conjugates revealed ubiquitylation also of OMM subdomains (**Fig. 3a**). Moreover, while not captured in the quantification of intra-mitochondrial signals (**Fig. 2i**), GFP-XIAP notably concentrated at OMMs prior to its accumulation within mitochondria (**Fig. 2h, Supplementary Movie 2**), in line with a role in acting upstream of Rab5 recruitment to valinomycin-depolarized mitochondria.

**Fig. 3:**
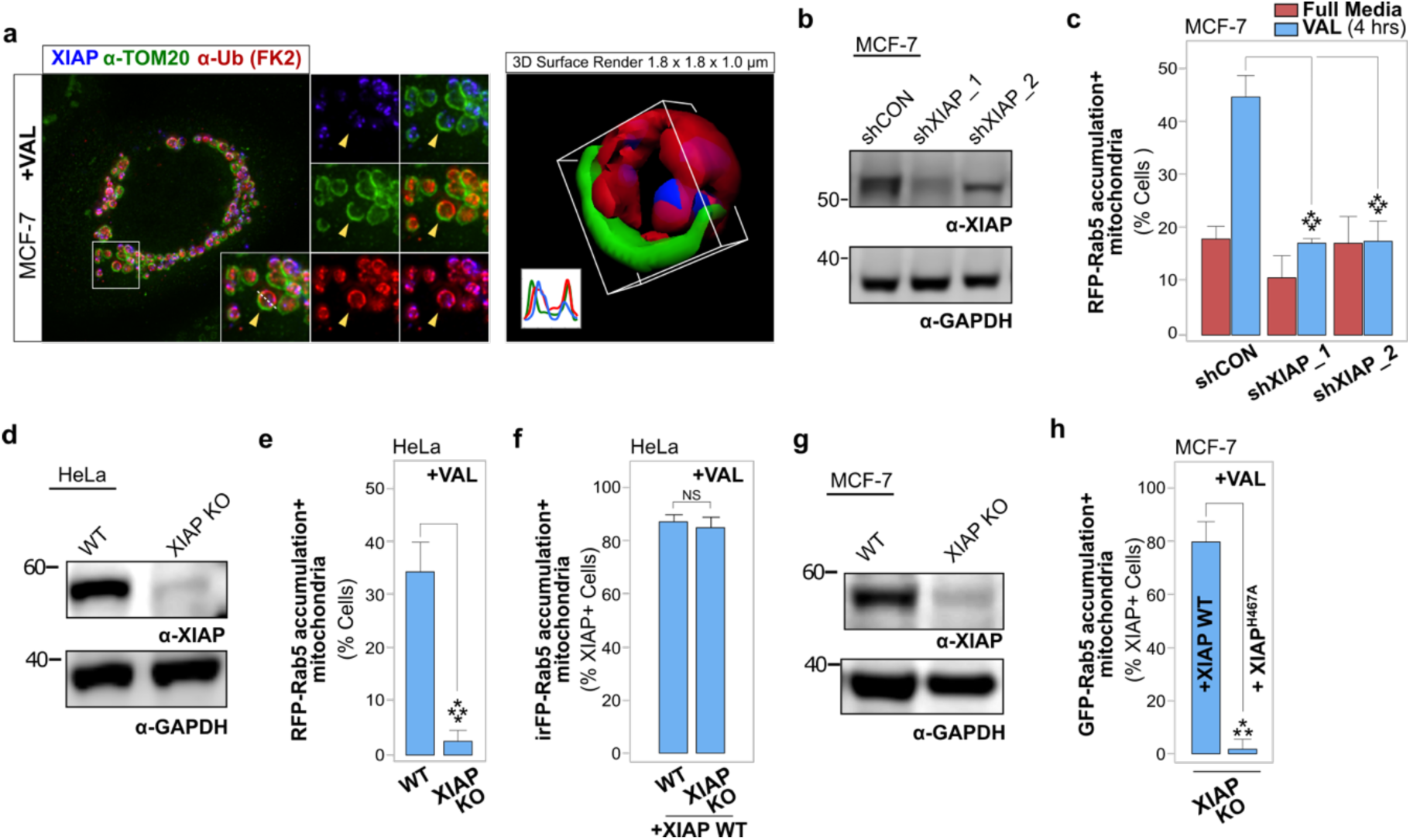
XIAP mediates endosomal targeting of stressed mitochondria, in a RING domain-dependent manner. **a** MCF-7 cell expressing RFP-XIAP, treated with VAL for 3 h. IF of TOM20 and of FK2 antibody-detected mono- and poly-ubiquitin conjugates. *Left*, overview and zoom images. *Right*, 3D surface rendering and plot profile of mitochondrion marked with yellow arrowhead in zoom images. **b** Representative immunoblot of XIAP protein levels in MCF-7 cells stably expressing shCON, shXIAP_1 or shXIAP_2. GAPDH, loading control. **c** MCF-7_shCON, MCF-7_shXIAP_1 and MCF-7_shXIAP_2 cells, co-expressing RFP-Rab5 and GFP-OMM, scored for RFP-Rab5+ mitochondria at 3 h VAL. Graph, mean ± SD of n = 3 experiments. ≥ 300 cells per condition. \*\*\*\**p* ≤ 0.0001 **d** Representative immunoblot of XIAP levels in HeLa WT and CRISPR/Cas9 XIAP knockout (KO) cells. GAPDH, loading control. **e** HeLa WT and HeLa XIAP KO cells, co-expressing RFP-Rab5 and GFP-OMM, scored for RFP-Rab5+ mitochondria at 3 h VAL. Graph, mean ± SD of n = 3 experiments. ≥ 120 cells per condition. \*\*\*\**p* ≤ 0.0001 **f** HeLa WT and HeLa XIAP KO cells co-expressing GFP-Rab5, iRFP-OMM and RFP-XIAP, scored for iRFP-Rab5+ mitochondria at 3 h VAL. Graph, mean ± SD of n = 3 experiments. ≥ 85 cells per condition. NS, not significant **g** Representative immunoblot of XIAP levels in MCF-7 WT and CRISPR/Cas9 XIAP KO cells. GAPDH, loading control. **h** MCF-7 WT and XIAP KO cells co-expressing GFP-Rab5 and RFP-XIAP or RFP-XIAP^H467A^, scored for GFP-Rab5+ mitochondria at 3 h VAL. Graph, mean ± SD of n = 3 experiments. > 130 cells per condition. \*\*\**p* ≤ 0.001 Scale bars, 10 µm.

Indeed, shRNA-mediated knockdown of XIAP (**Fig. 3b**) significantly reduced the valinomycin-activated mitochondrial Rab5 accumulation (**Fig. 3c**), identifying a role for endogenous XIAP in the rapid targeting of valinomycin-depolarized mitochondria by Rab5+ endosomes. Consistently, Rab5 targeting of depolarized mitochondria was abolished in HeLa XIAP knockout (KO) cells generated using CRISPR/Cas9 genome editing and reestablished by XIAP overexpression (**Fig. 3d-f**). Furthermore, in MCF-7 CRISPR/Cas9 XIAP KO cells, reconstitution with RFP-tagged wild type XIAP, but not with XIAP E3 ligase mutant XIAP^H467A 34^, reestablished mitochondrial Rab5 targeting (**Fig. 3g, h**), thus evidencing functional significance of XIAP-mediated ubiquitylation in endosome-mitochondrial interactions.

Together, these data substantiate a non-canonical role for XIAP ubiquitin E3 ligase as a central mediator of stress-induced endosome-mitochondrial interactions. Translocation of XIAP to OMMs preceded its intra-mitochondrial accumulation. Intra-mitochondrial accumulation of XIAP and Rab5 temporally coincided, which may indicate joint mitochondrial internalization. The presence of OMM ubiquitylation and reestablishment of mitochondrial targeting by Rab5 in XIAP depleted cells by reconstitution with wild type but not E3 ligase mutant XIAP, corroborate a role for XIAP-mediated ubiquitylation activity upstream of endosomal targeting to depolarized mitochondria. Moreover, the detection of intra-mitochondrial K63-linked polyubiquitin chains, subsequent to mitochondrial XIAP internalization, is suggestive of intra-mitochondrial XIAP E3 ligase activity.

### Endosomes target depolarized mitochondria independently of autophagy and are marked by distinct endocytic effectors and adaptors

To characterize endosomal vesicle signatures, we then investigated the presence of specific Rab5 effectors and endocytic adaptors on mitochondria-targeting endosomes. The Rab5 effectors APPL1 and EEA1 mark separate Rab5+ early endosomal subpopulations ^35^. Interestingly, RFP-APPL1+ vesicular structures were prominently recruited to XIAP+/Rab5+ mitochondria following valinomycin treatment (**Fig. 4a**). In contrast, valinomycin treatment did not result in RFP-EEA1 translocation to XIAP-targeted mitochondria (**Fig. 4b**). Instead, EEA1 remained with the cytosolic fraction of Rab5+ vesicles. We further tested whether valinomycin induces translocation of endocytic adaptors to the mitochondrial compartment. We found that OPTN, TAX1BP1 and, most prominently, Tollip, all relocated to depolarized mitochondria that had accumulated Rab5 and XIAP (**Fig. 4c-e**). In contrast, Parkin-mediated mitophagy in Parkin overexpressing cells engaged the recruitment of OPTN and TAX1BP1, but not Tollip (**Fig. 4f**). Importantly, the observed direct endosomal targeting of depolarized mitochondria did not involve macroautophagy. RFP-LC3 labeled autophagosomes displayed only limited transient interactions, with a minority of GFP-XIAP targeted mitochondria (**Fig. 5a, b, Supplementary Movie 4**). Moreover, while shRNA-mediated knockdown of Atg5 or Beclin1 (**Fig. 5c**), or CRISPR/Cas9-mediated deletion of autophagy-related Atg5 (**Fig. 5e, f**), effectively impaired LC3-monitored autophagy flux, it did not decrease Rab5-targeting to valinomycin-depolarized mitochondria (**Fig. 5d, g** and **h**).

**Fig. 4:**
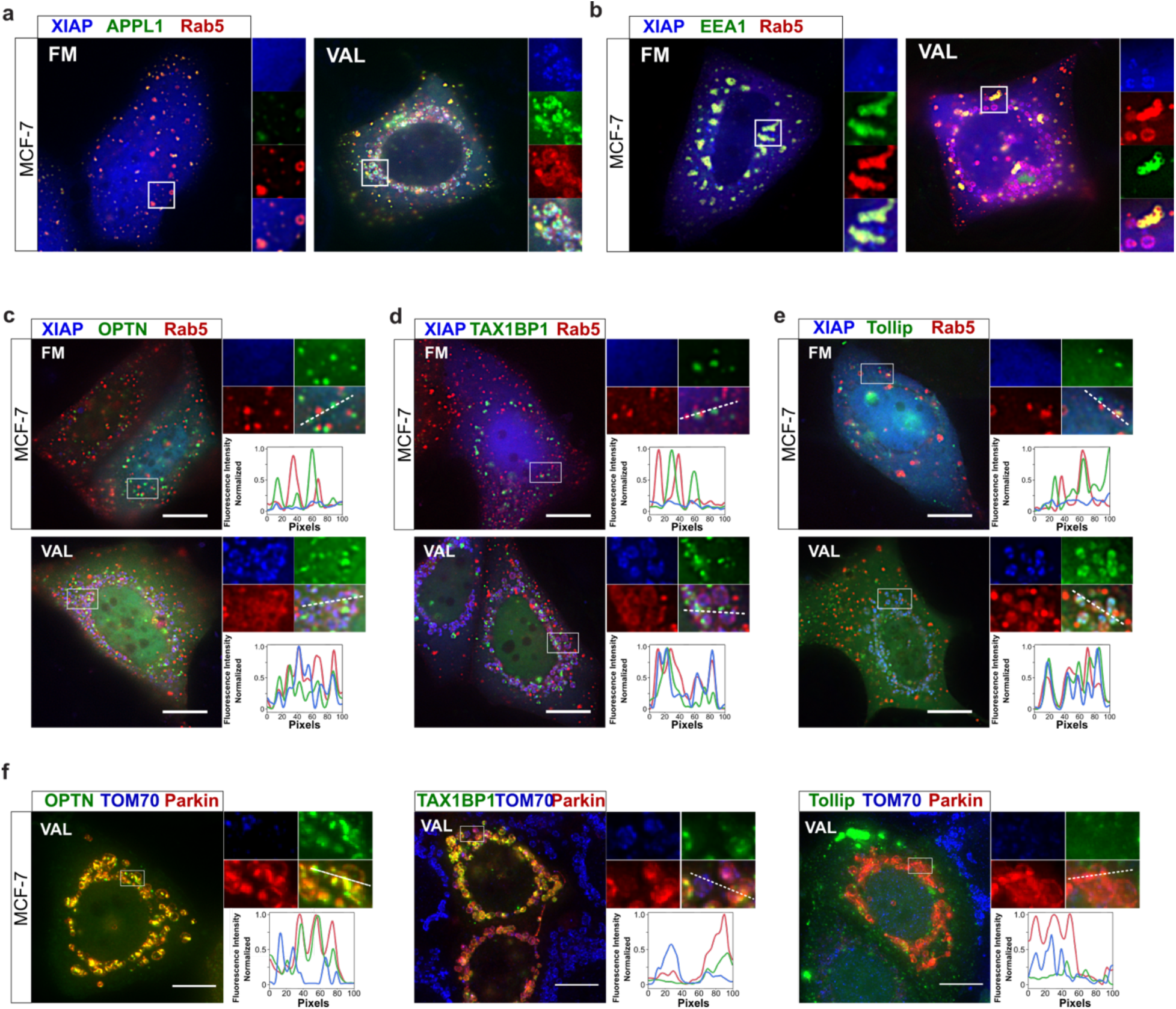
Endosomes targeting stressed, XIAP+ mitochondria are decorated with Rab5 effector APPL1 and endocytic adaptors OPTN, TAX1BP1 and Tollip. **a** MCF-7 cells co-expressing RFP-APPL1, GFP-XIAP and irFP-Rab5, imaged *live* in FM or at 3 h VAL. **b** MCF-7 cells co-expressing RFP-EEA1, GFP-XIAP and irFP-Rab5, imaged *live* in FM or at 3 h VAL. **c** MCF-7 cells co-expressing RFP-XIAP and irFP-Rab5 together with GFP-OPTN, imaged in FM or at 3 h VAL. Overview and ROI zoom images. Intensity profile over line in merged ROI. **d** MCF-7 cells co-expressing RFP-XIAP and irFP-Rab5 together with GFP-TAX1BP1, treated and presented as in (c). **e** MCF-7 cells co-expressing RFP-XIAP and irFP-Rab5 together with GFP-Tollip, treated and presented as in (c). **f** MCF-7 cells co-expressing RFP-Parkin with endocytic adaptors GFP-OPTN, GFP-TAX1BP1 or GFP-Tollip. IF of TOM70 at 3 h VAL. Scale bars, 10 µm.

**Fig. 5:**
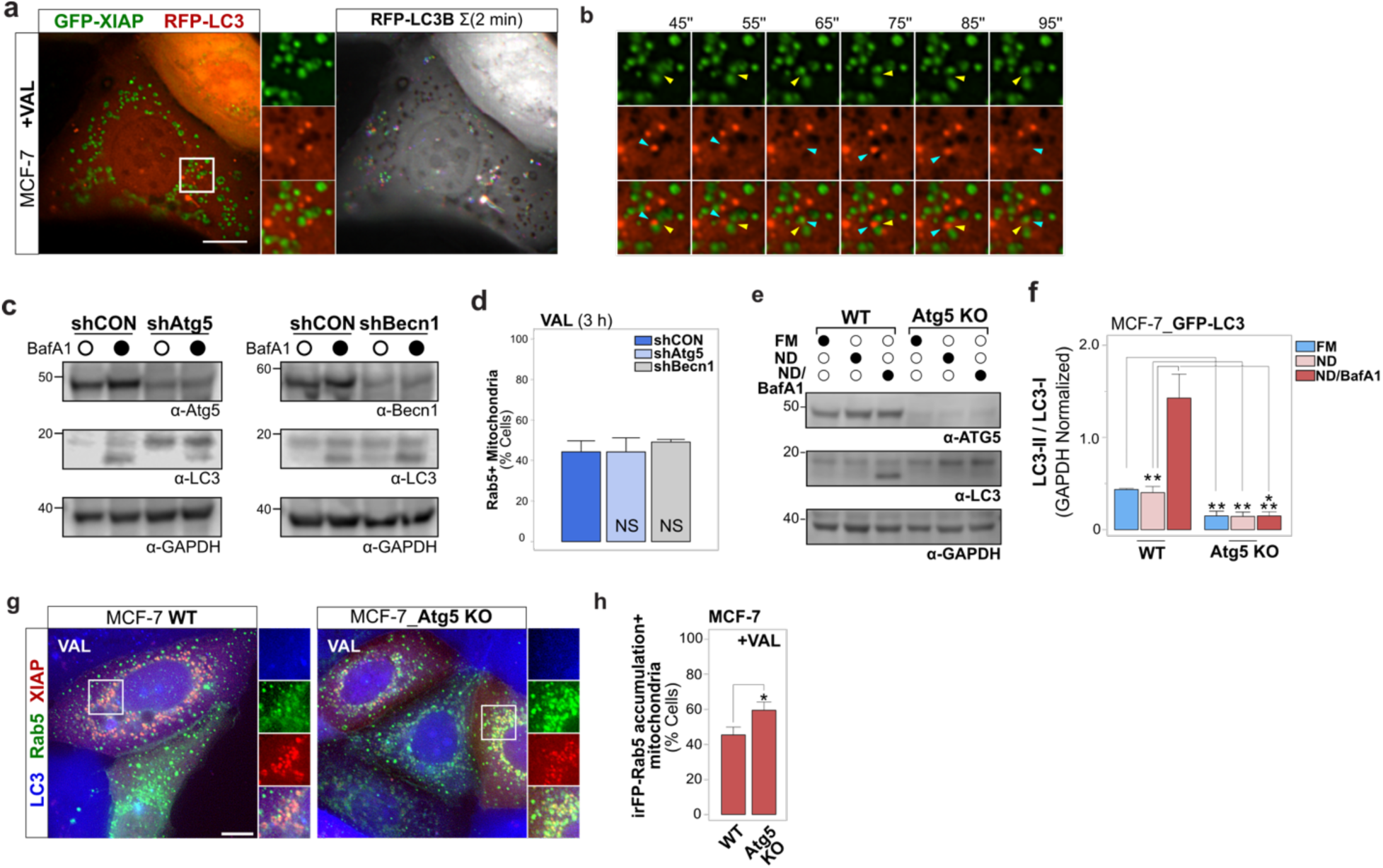
XIAP-mediated endosomal targeting of depolarized mitochondria operates independently of macroautophagy. **a** MCF-7 cells co-expressing GFP-XIAP and autophagosome marker RFP-LC3, *live* imaged every 5 s for 2 min at 3 h VAL. Single time-point color-merged overview image with single color and merged ROI zoom images, and 2-min time-projected image of RFP-LC3 movement (Σ2 min), color-coded by time-step. Note, mitochondria appear as voids within cytosolic RFP-LC3 signal. See also **Supplementary Movie 4**. **b** Time-series single color and merged zoom images of ROI from (a). Yellow arrowheads track exemplary mitochondrial-localized GFP-XIAP. Cyan arrowheads track an exemplary RFP-LC3+ autophagosome in transient vicinity to mitochondrion, without merging or engulfment. **c** Representative immunoblots of MCF-7 cells stably expressing shCON, shAtg5, or shBecn1, untreated or treated with BafA1 for 3 h. Detection of Atg5 or Beclin1, and LC3B-I and LC3B-II, and loading control GAPDH. **d** MCF-7_shCON, MCF-7_shAtg5 or MCF-7_shBecn1 cells transfected with RFP-Rab5 and subjected to 3 h VAL, scored for Rab5 accumulation in mitochondrial compartment (Rab5+ Mitochondria). Graph, mean ± SD of n = 3 experiments. ≥ 120 cells analyzed per condition. NS, not significant **e** Representative immunoblot of MCF-7_GFP-LC3 or MCF-7_GFP-LC3 CRISPR/Cas9 Atg5 KO cells, kept in FM, or subjected to nutrient deprivation (ND) for 3 h, without or with Bafilomycin A1 (BafA1). Immunodetection of LC3B-I and LC3B-II, ATG5, and loading control GAPDH. **f** Quantification of experiment in (e), quantifying autophagy flux (LC3-II / LC3-I). Graph, mean ± SD of n = 3 experiments. \**p* ≤ 0.05 **g** MCF-7_GFP-LC3 and MCF-7_GFP-LC3 Atg5 KO cells co-expressing RFP-XIAP and irFP-Rab5, in FM or at 4 h VAL. **h** Quantification of experiment in (g), scored for irFP-Rab5+ mitochondria at 3 h VAL. Graph, mean ± SD of n = 3 experiments. ≥ 200 cells analyzed per condition. NS, not significant Scale bars in fluorescence images, 10 µm.

Taken together, Rab5+ endosomes target depolarized mitochondria independently of macroautophagy, and are marked by specific endocytic effectors and adaptors. The selective presence of Rab5 effector APPL1, but not EEA1, on mitochondria-targeting early endosomes demonstrates targeting specificity and suggests relevance for APPL1-mediated signaling events ^36^. Notably, mitochondrial recruitment of Tollip represents a unique characteristic of the XIAP-mediated mitochondrial stress response, as Tollip does not participate in Parkin-mediated mitophagy ^37^.

### XIAP-mediated mitochondrial targeting by endolysosomes results in degradative processing of dysfunctional mitochondria

The above observed valinomycin-induced presence of K48- and K63-linked polyubiquitin chains and endosomes within inner mitochondrial compartments is suggestive of endolysosome-mediated degradative processing events. To gain insight into mitochondrial interactions with endolysosomal vesicles and resulting mitochondrial alterations at the ultrastructural level, we employed transmission electron microscopy. The overall population of control cell mitochondria exhibited their typical ultrastructure (**Supplementary Fig. 1a**) and interactions with electron-dense endolysosomal vesicles were detectable in only few instances (1%), in accordance with the above-described transient nature of endolysosome-mitochondrial interactions under normal growth conditions. In contrast, valinomycin-treated cells displayed abundant events of endolysosome-mitochondrial interactions (**Fig. 6a, Supplementary Fig. 1b**). Remarkably, detectable phenotypes ranged from near associations between endolysosomal vesicles and mitochondria, to vesicle-containing endocytic cup-like invaginations of mitochondrial membranes, to vesicles surrounded by double-membrane structures inside mitochondria, together with varying portions of cytoplasm. For the latter phenotype, it is not possible to discriminate between the presence of vesicles within deep mitochondrial membrane invaginations ^38^ *vs*. mitochondrial internalization of membrane-enclosed vesicles. Nevertheless, some mitochondria contained several vesicles of various electron-densities and substructures, free from mitochondrial membrane enclosure, supporting uptake into inner mitochondrial compartments. Of note, the ultrastructure of these mitochondria appeared severely altered, with merely cristae remnants, suggestive of substantial intra-mitochondrial remodeling and degradation.

**Fig. 6:**
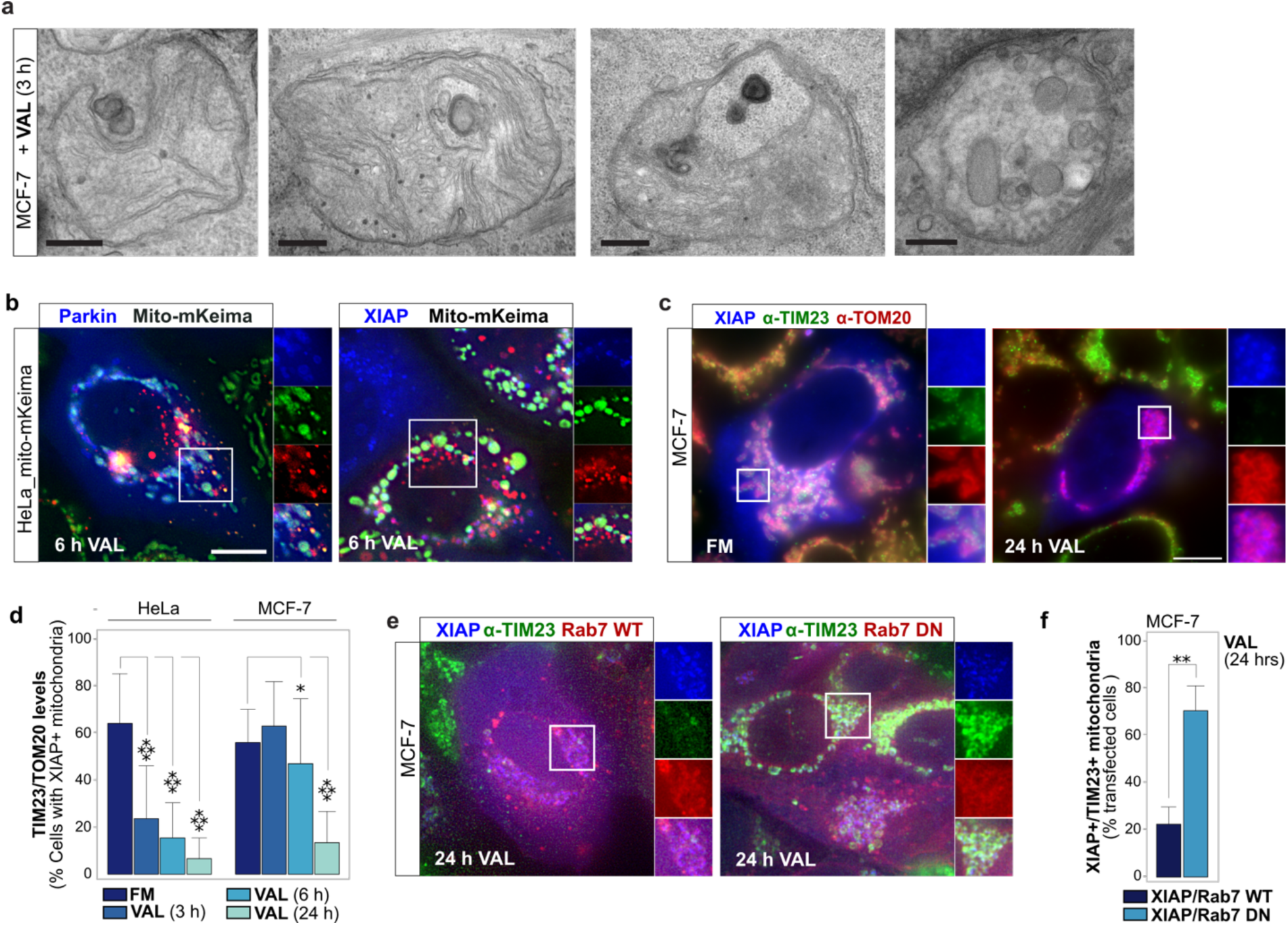
XIAP mediates sub-organelle level degradative processing of stressed mitochondria. **a** Transmission electron micrographs of MCF-7 cells treated with VAL for 3 h. Scale bars, 200 nm. See also **Supplementary Fig. 1**. **b** HeLa cells co-expressing mito-mKeima and GFP-Parkin or GFP-XIAP at 6 h VAL. Representative examples of GFP signal (blue), neutral pH mito-mKeima (green), low pH lysosomal mito-mKeima (red). **c** MCF-7 cells expressing GFP-XIAP. IF of TIM23 and TOM20 at 24 h FM or VAL. Merged color overview images and single and merged color zoom images of ROI. **d** MCF-7 and HeLa cells expressing RFP-XIAP, in FM or treated with VAL for 3, 6 or 24 h. IF of TIM23 and TOM20. Graph, mean ± SD for single cell TIM23/TOM20 ratios, ≥ 15 cells per condition, n = 2 experiments. \**p* ≤ 0.05, \*\*\*\**p* ≤ 0.0001 **e** MCF-7 cells co-expressing GFP-XIAP and RFP-Rab7 WT or dominant negative (DN) RFP-Rab7^T22N^. IF of TIM23 at 24 h VAL. Merged color overview images and single and merged color zoom images of ROI. **f** Quantifications of TIM23 levels in cells as in (e). Graph, mean ± SD of n = 3 experiments. ≥ 250 cells per condition. \*\**p* ≤ 0.01 Scale bars, 10 µm.

To assess degradative endolysosomal processing of mitochondria we utilized mitochondrial matrix-targeted mito-mKeima, which is designed to detect lysosomal degradation during mitophagy based on lysosomal acidification-induced spectral shifts of mito-mKeima fluorescence ^37,39^. Similarly to as observed with Parkin-mediated mitophagy in response to valinomycin treatment, XIAP-targeted valinomycin-depolarized mitochondria were positive for acidic pH-induced red mito-mKeima fluorescence (**Fig. 6b**), suggesting uptake of mito-mKeima+ mitochondrial matrix components into endolysosomes. In addition, we quantified cellular levels of inner mitochondrial membrane (IMM) protein TIM23, relative to those of OMM protein TOM20, in GFP-XIAP transfected cells. Cellular TIM23 levels were significantly decreased relative to TOM20, by 6 h in MCF-7 cells and 3 h in HeLa cells (**Fig. 6c, d**). Notably, the late endosomal and lysosomal GTPase Rab7 also colocalized with mitochondria in response to valinomycin treatment, in line with mitochondrial targeting by degradation-competent endolysosomal vesicles (**Fig. 6e**). Importantly, interference with endolysosomal function through expression of the dominant-negative mutant Rab7^T22N^ ^40^ impaired TIM23 degradation (**Fig. 6e, f**).

The above findings support a link between XIAP-mediated mitochondrial ubiquitylation and endolysosome-mediated degradation of inner mitochondrial compartment components.

### XIAP-mediated mitochondrial processing competes with Parkin-dependent mitophagy

XIAP-mediated targeting of endocytic machinery to apoptotic or depolarized mitochondria does not require Parkin ^23^ and, differing from Parkin-mediated mitophagy (**Fig. 4f**) ^37^, it involves the recruitment of endocytic adaptors including Tollip (**Fig. 4c-e**). In the following, we addressed whether the mitochondrial processing modes activated by XIAP vs. Parkin E3 ligases undergo competitive regulation.

First, we addressed a potential regulatory role for PINK1, a serine/threonine kinase and upstream regulator of Parkin-mediated mitophagy ^37^. N-terminal RFP-tagging of PINK1 creates a constitutively active form of PINK1 ^41^. As reported, expression of RFP-PINK1^CA^ was sufficient to translocate GFP-Parkin to mitochondria in MCF-7 cells, but did not trigger GFP-XIAP translocation (**Fig. 7a-c**). Moreover, valinomycin-triggered mitochondrial translocation of GFP-XIAP was unaffected by RFP-PINK1^CA^ co-expression (**Fig. 7b, c**). We conclude that PINK1 neither functions as an upstream initiator, nor negative regulator, of XIAP-mediated mitochondrial processing.

**Fig. 7:**
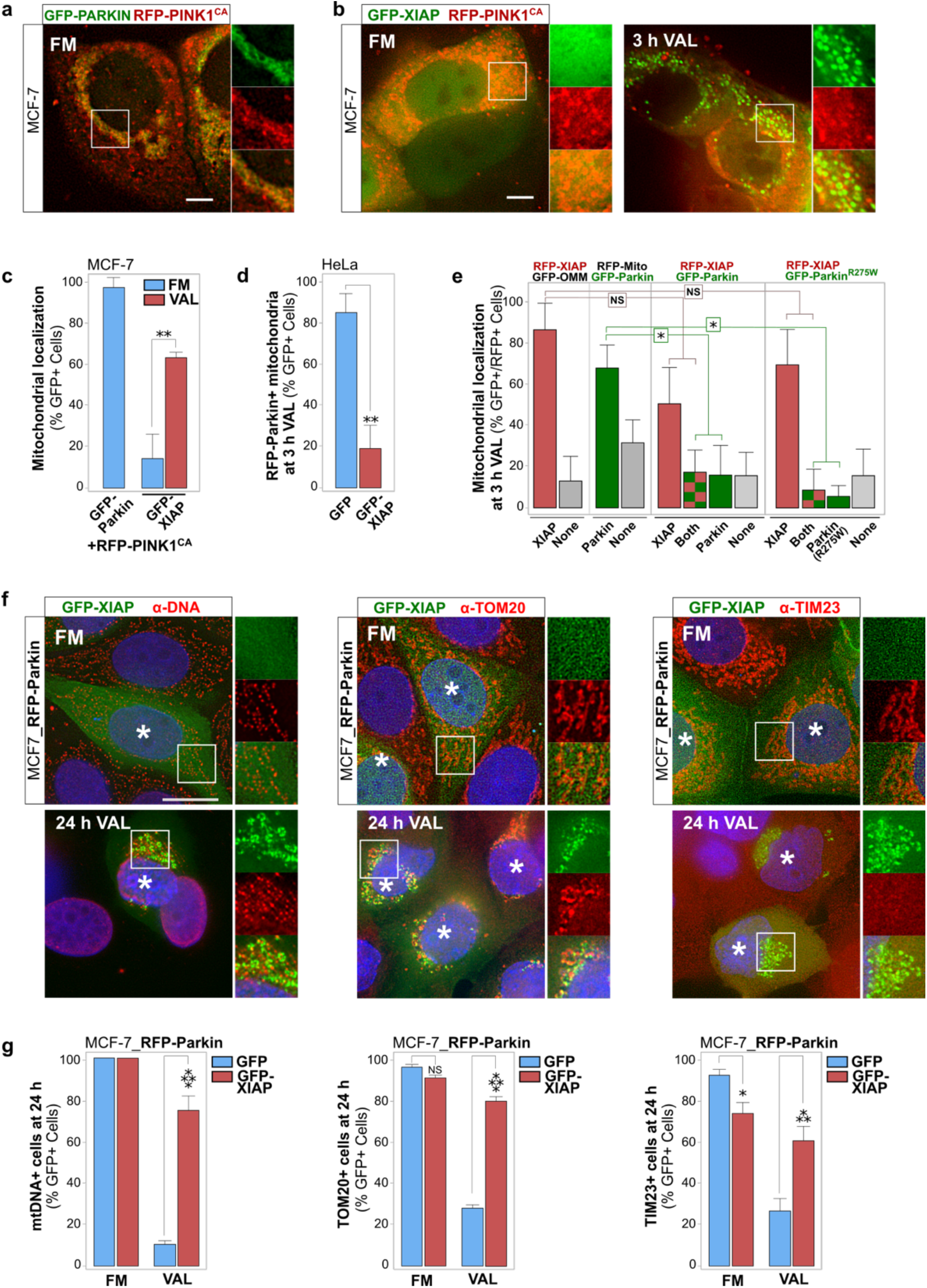
Ubiquitin E3 ligases XIAP and Parkin exert distinct, competitive modes of mitochondrial processing. **a** MCF-7 cells co-expressing GFP-Parkin and constitutively active RFP-PINK1 (RFP-PINK1^CA^), imaged *live* in FM. Merged color overview, and single and merged color ROI zoom images. **b** MCF-7 cells co-expressing GFP-XIAP and RFP-PINK1^CA^, imaged *live* in FM or at 3 h VAL. **c** Quantification of experiments in (a) and (b). Cells scored for mitochondrial translocation of GFP-Parkin or GFP-XIAP. Graph, mean ± SD of n = 3. ≥ 220 cells per condition. \*\**p* ≤ 0.01 **d** HeLa cells co-transfected with RFP-Parkin and GFP or GFP-XIAP. Cells scored for mitochondrial localization of RFP-Parkin at 3 h VAL. Graph, mean ± SD of n = 3. ≥ 100 cells per condition. \*\**p* ≤ 0.01 **e** MCF-7 cells co-transfected with RFP-XIAP and GFP-Parkin together or with indicated mitochondrial marker, or E3 ligase mutant GFP-Parkin^R275W^. Cells scored for mitochondrial localization of RFP-XIAP and/or GFP-Parkin or GFP-Parkin^R275W^ at 3 h VAL. Graph, mean ± SD of n = 3. ≥ 150 cells per condition. \**p* ≤ 0.05 **f** MCF-7 cells stably expressing RFP-Parkin, transfected with GFP-XIAP (marked with a white asterisk). IF of mitochondrial DNA, TOM20 or TIM23 at 24 h of FM or VAL. **g** MCF-7 cells stably expressing RFP-Parkin, transfected with GFP or GFP-XIAP. IF of mitochondrial DNA, TOM20 or TIM23 at 24 h of FM or VAL. Cells scored for mitochondrial presence of indicated IF signals. Graph, mean ± SD of n = 3. > 260 cells per condition. *p < 0.05, ***p < 0.001, ****p < 0.0001

Second, we determined whether XIAP and Parkin E3 ligases undergo competitive regulation in responding to mitochondrial depolarization. Indeed, GFP-XIAP expression in HeLa cells stably expressing RFP-Parkin blocked mitochondrial translocation of RFP-Parkin in response to valinomycin (**Fig. 7d**). Valinomycin treatment of MCF-7 cells co-expressing a mitochondrial marker with either RFP-XIAP or GFP-Parkin, or co-expressing both E3 ligases, confirmed a competitive nature of mitochondrial targeting also in MCF-7 cells (**Fig. 7e**). At 3 h of treatment in the majority of cells either RFP-XIAP or GFP-Parkin were recruited to mitochondria, while only a small percentage of cells displayed co-recruited RFP-XIAP and GFP-Parkin. Importantly, similar to as in HeLa cells, increasing XIAP protein levels through RFP-XIAP expression largely inhibited GFP-Parkin translocation to mitochondria. In contrast, co-expression of GFP-Parkin reduced RFP-XIAP translocation by a substantially lesser extent. Parkin interference with XIAP translocation was dependent on Parkin E3 ligase activity, as the Parkin E3 ligase mutant GFP-Parkin^R275W^ ^42^ did not impact translocation of RFP-XIAP.

To explore XIAP vs. Parkin pathway competition on the functional level of mitochondrial degradation, we expressed GFP or GFP-XIAP in MCF-7 cells stably expressing RFP-Parkin and analyzed the degradation of the mitochondrial membrane proteins TOM20, TIM23 and mitochondrial DNA (mtDNA) in cells expressing RFP-Parkin alone, vs. in cells co-expressing RFP-Parkin and GFP or GFP-XIAP (**Fig. 7f, g**). As expected, in cells expressing RFP-Parkin alone, or in combination with GFP, 24 h of valinomycin treatment resulted in clearance of mtDNA, and TOM20 and TIM23. In contrast, cells co-expressing RFP-Parkin and GFP-XIAP had maintained mtDNA and OMM-localized TOM20 at 24 h of valinomycin, indicating that XIAP exerts an inhibitory function over Parkin-mediated mitophagy, potentially influenced by the relative cellular abundance of each of the two E3 ligases. Consistent with our above findings (**Fig. 6c**), IMM-localized TIM23 was degraded also in XIAP co-expressing cells.

In summary, these data demonstrate that XIAP and PINK1/Parkin mediate distinct and competitive mitochondrial processing modes. Notably, co-expression experiments suggest that XIAP negatively regulates Parkin mitophagy in a concentration-dependent manner, at the levels of both Parkin translocation and mitochondrial degradation.

## Discussion

Inter-organelle communication is an increasingly recognized fundamental component of cellular function ^7^. Notably, questions concerning whether, when and how endolysosomes directly interact with mitochondria have only recently begun to be addressed ^16^. Here, we implemented a sensitive *live* microscopy approach to quantitatively describe interactions between endolysosomes and mitochondria under energetic and oxidative stress conditions. We describe a pathway for largescale, switch-like targeting of endolysosomes to stressed mitochondria. We show that this pathway can function as an autophagy-independent intra-mitochondrial processing mode, that is mediated by XIAP E3 ligase and associated with routing of endocytic adaptor- and effector-marked endolysosomal vesicles to mitochondria.

Intriguingly, our electron microscopy and high-resolution fluorescence microscopy studies together suggest that endolysosomal vesicles enter inner mitochondrial compartments. This would differ remarkably from other vesicle-based mitochondrial quality control programs. For example, canonical mitophagy ^13^, and the recently proposed direct uptake of mitochondria into lysosomes ^43,44^ or early endosomes ^27^, are all characterized by mitochondrial engulfment into degradative organelles. The here described pathway is also distinct from the process of mitochondrial-derived vesicles (MDVs) ^14^.

In addition to these mechanistic distinctions, our dynamics analyses show that XIAP-mediated mitochondrial processing acts rapidly. At about 3 h following depolarization Rab5+ endosomes target the entire cellular mitochondrial population, in a switch-like manner that is completed within minutes once triggered (**Supplementary Movie 1**). By comparison, at 4 h following mitochondrial uncoupling ∼4-15 MDVs ^45^ or ∼10 early endosome-engulfed mitochondria ^27^ are observed per cell, and Parkin-mediated mitophagy leads to gross mitochondrial degradation on a time-scale of typically 12-48 h ^46^. Also, (macro-)autophagy independence eliminates kinetic and energetic costs of autophagosomal membrane biogenesis cascades. The re-routing of existing organelles and post-translational modifications through XIAP E3 ligase might in part explain the rapid nature of the here-described pathway. Of note, recent reports describe stress-responsive degradative processing of the ER organelle compartment through direct degradation of ER subcomponents by endolysosomes, independent of (macro-)autophagy ^47,48^, and a proteomics-based study indicates the importance of autophagy-independent mechanisms in mitochondrial protein turnover ^49^. We propose that XIAP-mediated mitochondrial processing is particularly suited for swift response to acute stressors. In addition, our data suggest that this pathway might contribute also to homeostatic mitochondrial quality control, as we detected isolated instances of mitochondria-targeted vesicles under normal growth conditions.

Under oxidative stress conditions, the presence of endolysosomal vesicles in mitochondria was associated with profound remodeling of inner mitochondrial compartments. Although it is at present mechanistically unclear how internalized endolysosomes may accomplish degradation, our electron micrographs clearly demonstrate substantial loss of cristae. Consistent with this, and similar to our previous findings during tBid-mediated apoptosis signaling ^23^, we observed ubiquitylation inside OMM-enclosed compartments. Moreover, our data support the exposure of mitochondrial matrix-localized mt-mKeima to the acidic pH of endolysosomes and identify IMM-resident TIM23 as a protein targeted for degradation, in a Rab7-dependent manner. Quantitative mass spectrometry-based approaches will be instrumental to comprehensively determine the select mitochondrial components that are subject to XIAP-mediated ubiquitylation and/or degradation.

In addition to functioning as a mitochondrial processing pathway, we speculate that the merging of mitochondrial and endolysosomal organelle compartments creates a distinct, spatially compartmentalized signaling platform. Clues into particular signaling functions may be contained in the pathway participation of specific APPL1+/EEA1– endosomal subpopulations, and distinctive endocytic regulators and adaptors. XIAP engages endocytic adaptor subsets distinct from Parkin. In addition to OPTN and TAX1BP1, both also participating in Parkin-mediated mitophagy ^37^, XIAP uniquely targets Tollip to stressed mitochondria. Notably, Tollip participates in the regulation of inflammatory signaling ^50,51^. Furthermore, the here described valinomycin-induced pathway differs from the recently reported endosome-mitochondrial interactions during high-level oxidative stress induced by laser irradiation or 250 µM H_2_O_2_, in which Rabenosyn-5+/EEA1–/APPL1– endosomal subpopulations were found to target mitochondria ^24^. This prompts the hypothesis that there exist multiple, stress-specific modes of mitochondrial targeting by specific endosomal subpopulations, that are orchestrated by distinct endocytic regulators.

Importantly, XIAP is ubiquitously expressed in human tissues and its expression is elevated in various cancers ^52^, and we found that endogenous XIAP levels are sufficient for pathway operation. Conversely, Parkin mitophagy research models typically employ exogenous overexpression, as Parkin is commonly deleted in human cancers ^53^ and is weakly expressed in most tissues (**Supplementary Fig. S2**). Co-expression of XIAP and Parkin in MCF-7 breast cancer cells revealed that XIAP-mediated mitochondrial processing is not only distinct from, but also can outcompete Parkin-mediated mitophagy. Interestingly, both mitochondrial XIAP and Parkin pathways are under the control of BCL-2 family protein signaling. Parkin-mediated mitophagy is negatively regulated by anti-apoptotic BCL-2 proteins and does not require pro-apoptotic BAX ^54^, and Parkin targets BAX for degradation ^55^. We found that XIAP targeting of stressed mitochondria is also suppressed by BCL-X_L_, but necessitates presence of BAX or BAK ^32^. Future efforts will establish mechanistic insights into the nature of competition between the mitochondrial actions of XIAP vs. Parkin E3 ligases, including a competition for the supply of E1 activated-ubiquitin, and E2 ubiquitin-conjugating enzymes such as UbcH5 that are utilized by both XIAP ^56^ and Parkin ^57^. Moreover, it will be important to further our understanding of a potential direct impact of XIAP on autophagy regulation ^58-60^ and investigate the extent of its contribution on pathway engagement.

Of note, we recently reported that during drug-induced intrinsic apoptosis, endolysosomal targeting of mitochondria contributes to BAX-mediated mitochondrial permeabilization and cell death ^61^. Consequently, our identification of XIAP as an upstream mediator of endolysosome-mitochondrial interactions suggests a pro-MOMP role for XIAP during apoptosis signaling. Indeed, we and others have shown that XIAP can positively regulate MOMP activation, while counteracting post-MOMP caspase-mediated cell death signaling, including via the initiation of Smac/DIABLO degradation through concerted endolysosomal and proteasomal action ^23,32,62^. These findings indicate that XIAP exerts a multi-layered control over both induction and functional outcomes of MOMP, and suggest that XIAP’s role in cell death regulation is more complex than previously appreciated. It is tempting to speculate that specific modes of endolysosome-mitochondrial interactions partake in the fine-tuning of pro-survival vs. pro-death mitochondrial responses to sub-lethal vs. lethal stress. Future work will be needed to dissect the stress-specific live/death regulation through endolysosome-mitochondrial interactions and address the complexity of integration by core regulators of apoptosis signaling, including XIAP.

In summary, our data elucidate the spatio-temporal dynamics and molecular characteristics of an oxidative stress-induced mitochondrial processing mode that is distinct from canonical programs and furthers our insights into the interconnectivity between mitochondrial and endolysosomal organelle compartments under oxidative stress conditions. Our work proposes new directions in mitochondrial signaling and highlights the need for integrative understanding of organelle biology in health and disease.

## Materials and Methods

### Cell culture and transient transfections

Human MCF-7 breast cancer (CLS Cell Lines Service) and HeLa cervical cancer (ATCC® CCL-2™), were maintained in DMEM (1 g/L D-glucose, 0.11 g/L sodium pyruvate), the Lenti-X embryonic kidney 293T cell line (Clontech) in DMEM (4.5 g/L D-glucose). Media were supplemented with 10% FBS (Sigma), GlutaMAX (Gibco), non-essential amino acids (Gibco), penicillin/streptomycin/amphotericin B (Gibco) and 100 μg/mL Normocin (Invivogen). Cell lines were routinely controlled for mycoplasma contamination using Hoechst 33342 (1 µg/mL). Transient transfections were performed using jetPRIME (Polyplus) and culturing media renewed 4-6 h following transfection. Experiments were performed following 16-24 h of expression.

### Stable cell line generation

Human MCF-7 breast cancer (CLS Cell Lines Service, no. 300273) and HeLa cervical cancer (ATCC® CCL-2™), were maintained in DMEM (1 g/L D-glucose, 0.11 g/L sodium pyruvate), the Lenti-X embryonic kidney 293T cell line (Clontech, no. 632180) in DMEM (4.5 g/L D-glucose). Media were supplemented with 10% FBS (Sigma, no. F2442), GlutaMAX (Gibco), non-essential amino acids (Gibco), penicillin/streptomycin/amphotericin B (Gibco) and 100 μg/mL Normocin (Invivogen). Cell lines were routinely controlled for mycoplasma contamination using Hoechst 33342 (1 µg/mL). Transient transfections were performed using jetPRIME (Polyplus) and culturing media renewed 4-6 h following transfection. Experiments were performed following 16-24 h of expression.

MCF-7 cell lines stably expressing GFP-LC3 (MCF-7_GFP-LC3) or RFP-Parkin (MCF-7_RFP-Parkin) were generated via transient transfection, followed by selection and expansion from single cell colonies with G418 (500 μg/mL; Invitrogen).

Lentiviral stable transduction was performed by transfecting psPAX2 (Addgene, no. 12260), pCMV-VSV-G (Addgene, no. 8454) and the respective pLKO.1 shRNA or lentiCRISPR v2 construct into Lenti-X 293T cells. Virus particle-containing supernatants were harvested and used to infect MCF-7 cells. Selection of stably shRNA-expressing polyclonal cell lines was achieved using puromycin (1 µg/mL; Gibco). pLKO.1 with non-targeting sequence 5’-AATTGCCAGCTGGTTCCATCA-3’ was used to generate shRNA control (shCON) cell lines. For lentiviral gene knockdown, pLKO.1 shRNA was directed against the following human target sequences:

5’-GACAGTTTGGCACAATCAATA-3’ (Beclin1),

5’-AGCTGTAGATAGATGGCAATA-3’ (XIAP #1)

5’-GCACTCCAACTTCTAATCAAA-3’ (XIAP #2).

For CRISPR/Cas9-mediated genome editing, MCF-7 or HeLa single cell colonies were selected from cells transduced with the following sequences targeting human genes cloned into lentiCRISPR v2 (Addgene no. 52961):

5’-AACTTGTTTCACGCTATATC-3’ (ATG5),

5’-TACAAGAAGCTATACGAATG-3’ (XIAP).

Stable cell lines were maintained in selection media, and cultured in selection-free medium for at least 24 h prior to experiments.

### Plasmids

The following fluorescent protein constructs were used as previously described: tagRFP-Parkin, EGFP-Rab5A, tagRFP-XIAP and EGFP-XIAP (all from ^23^); iRFP670-Rab5, GFP-OMM and iRFP670-OMM (all from ^61^); pEGFP-OPTN ^63^, pEGFP-TAX1BP1 ^64^, pQCXIP-Vx3K0-GFP

(Addgene; no. 35527 ^33^), EGFP-Rab7 and dominant negative mutant EGFP-Rab7^T22N^ ^40^, GFP-LC3B ^65^.

Fluorescent protein constructs generated for this study: PCR-amplified from human cDNA: tagRFP-PINK1, tagRFP-APPL1, tagRFP-EEA1, EGFP-Tollip. Generated through site-directed mutagenesis as previously described were E3 ligase-deficient tagRFP-XIAP^H467A^ ^34^, tagRFP-Parkin^R275W^ ^42^. To generate outer mitochondrial membrane (OMM) markers BFP-OMM, tagBFP was fused to the OMM-targeting transmembrane domain of Fis1 ^66^. Generated through subcloning: tagRFP-Rab5 from EGFP-Rab5 ^67^, EGFP-Parkin from tagRFP-Parkin ^67^. mt-mKeima (Addgene; no. 72342 ^37^) was sub-cloned into pLJM1-EGFP (Addgene; no. 19319 ^68^), replacing EGFP. In text and figure legends, tagRFP and mCherry are referred to as RFP, iRFP670 is referred to as irFP, EGFP as GFP, and tagBFP as BFP.

### Drug treatments and nutrient deprivation

Cells were treated with drugs diluted in phenol red-free fully supplemented media (FM; Gibco) at 37°C for the indicated durations at the following concentrations: bafilomycin A1 (BafA1; 0.1 µg/mL; Alfa Aesar), carbonyl cyanide m-chlorophenyl hydrazine (CCCP; 20 µM; EMD), hydrogen peroxide (H_2_O_2_, 250 µM; Sigma-Aldrich), nigericin (10 µM; Sigma-Aldrich), oligomycin/antimycin A (OA; 10 µM/25 µM; Cell Signaling; Sigma), Trolox (Tx; 200 µM; EMD Millipore); and valinomycin (VAL; 0.9 µM; Sigma). For nutrient deprivation (ND), cells were incubated in glucose containing HBSS (Invitrogen).

### Immunofluorescence

For fluorescence detection of endogenous proteins, cells were plated in 8-well microscopy μ-slides (Ibidi; no. 80826), transfected and treated as indicated, and fixed in electron microscopy-grade methanol-free paraformaldehyde (PFA, 4% in PBS; EMS) for 15 min. Following, cells were permeabilized with 0.3% Triton X-100 in PBS for 10 min and blocked with 3% BSA for 45 min. Cells were then incubated with primary antibodies against, DNA (American Research Products), Multiubiquitin Chains (Clone FK2; Cayman Chemical), TIM23 (BD Biosciences), TOM20 (Santa Cruz Biotechnology), TOM70 (Santa Cruz Biotechnology), ubiquitin, K48-linkage-specific (Merck Millipore), ubiquitin, K63-linkage-specific (Merck Millipore) for 1 h at room temperature, or with Rab5 (Cell Signaling), or XIAP (Santa Cruz Biotechnology) antibodies at 4°C overnight. Fluorescent staining was performed for 30 min at room temperature using highly cross-adsorbed Alexa Fluor 405, 488, 546 or 647 secondary antibodies (Invitrogen). Following primary and secondary antibody incubations, cells were washed three times in PBS. Where indicated, nuclei were stained with Hoechst 33342 (1 µg/mL) in PBS for 10 min.

### Fluorescent dye loading

To detect lipid peroxidation, cells were incubated with Bodipy 581/591 C11 (Thermo Fisher Scientific, no. D3861) at 1 µM for 1 h. Loading was performed at 37°C and 5% CO_2_ and followed by placing of cells in fresh media.

### Detection of mitochondrial processing using mito-mKeima

HeLa cells stably expressing mitochondrial matrix-targeted mKeima (mito-mKeima ^37,39^) were transfected with either GFP-XIAP or GFP-Parkin. Cells were imaged for GFP (blue excitation/green emission), neutral mKeima (blue excitation/red emission (neutral pH), and acidic mKeima (green excitation/red emission).

### Fluorescence imaging

Cells were plated in 8-well microscopy μ-slides (Ibidi), transfected and treated as indicated, and imaged *live* or following fixation in 4% PFA. Widefield fluorescence microscopy was performed with a DeltaVision Elite microscope system (GE Healthcare), equipped with a Scientific CMOS camera (Chip size: 2560 × 2160 pixels), an UltraFast solid-state illumination, an environmental chamber for *live* cell imaging at 37°C and 5% CO_2_, a 60x (N.A. 1.42) oil immersion objective, the UltimateFocus module, and 488 nm and 568 nm laser modules. Single image slices, 2D Z-stacks using optical axis integration (OAI), or Z-stacks using 0.2 μm step increments were acquired and deconvolved (Softworx, Applied Precision).

### Image analysis, quantification, and presentation

Image preparation and analysis was performed using Fiji ^69^. Image pseudocolors correspond to font colors of protein labels. 3D surface renderings were built from Z-stacks captured in 0.2 µm steps. Deconvolved slices were contrast adjusted in Fiji before implementation of the ImageJ 3D Viewer plugin. Blue TOM20 or TOM70 immunofluorescence signals are set to 30-40% transparency in 3D surface renderings displaying three channels.

Stack projections of *live* time series imaging were generated to represent 3D and 4D dynamics. Stacks of calculated inter-organelle interactions were combined, sum-projected, and pseudo-colored as indicated. Resulting images are sometimes merged with raw or filtered images of whole channels. Sum projection and pseudo-coloring of time-steps was performed using the ‘Image -> Hyperstacks -> Temporal-Color Code’ function. Alternatively, ROIs from combined stacks of mitochondria, endosomes, and interactions were re-ordered by swapping the X (space) and Z (time) axes using ‘Image -> Hyperstacks -> Re-Order Hyperstacks…’ in Fiji. The resulting YT-coordinate sum images demonstrate the stability of interactions over time.

Intensity profiles were measured using ‘Analyze -> Plot Profile’ on images contrast adjusted in Fiji. Fluorescence intensities were normalized and plotted in colors corresponding to displayed images. Movies were prepared from *live* cell time series images using Fiji. Contrast adjusted channels or interactions were merged as shown. Scale bars, and labels were generated and combined in Fiji.

### Ultrathin-section transmission electron microscopy (TEM)

MCF-7 cells transiently expressing GFP-Rab5, to assess pathway activation by fluorescence microscopy prior to fixation, were subjected to the indicated treatments and fixed in 2.5% glutaraldehyde (Electron Microscopy Sciences) in 0.1 M sodium cacodylate buffer (pH 7.4) for 1 h at room temperature, followed by three washes in 0.1 M cacodylate buffer. Subsequently, cells were post-fixed for 1 h in 1% osmium tetroxide (Electron Microscopy Sciences) in the same buffer at room temperature. Next, samples were washed in water and stained for 1 h at room temperature in 2% uranyl acetate (Electron Microscopy Sciences), and then washed again in water and dehydrated in a graded series of ethanol. The samples were then embedded in Embed-812 epoxy resin (Electron Microscopy Sciences). Ultrathin (50–60-nm) sections were cut using an Ultracut ultramicrotome (Reichart-Jung) and collected on formvar- and carbon-coated nickel grids, stained with 2% uranyl acetate and lead citrate before examination with a Philips/FEI BioTwin CM120 electron microscope under 80 kV. Mitochondria were identified based on a surrounding double-membrane, cristae or cristae remnants, and oftentimes contained calcium phosphate crystals.

### Western blotting

Whole-cell lysates were prepared with RIPA lysis buffer (Millipore) containing cOmplete EDTA free protease inhibitor cocktail (Roche). Protein concentrations were determined using Coomassie reagent (Sigma-Aldrich). Samples were denatured in LDS sample reducing buffer (Invitrogen), electrophoresed using Bis-Tris NuPAGE gels (Invitrogen) and proteins were transferred to nitrocellulose using the iBlot dry blotting system (Invitrogen). Immunodetection was performed using antibodies against Atg5 (Cell Signaling; no. 12994S), Beclin1 (Cell Signaling; no. 3495S), LC3 (Cell Signaling; no. 3868S), GAPDH (Santa Cruz Biotechnology; no. sc-25778), Rab5 (Cell Signaling; no. 3547S), XIAP (Santa Cruz Biotechnology; no. sc-55551). Horseradish peroxidase-conjugated secondary antibodies (Rockland) and home-made ECL substrate were used to visualize immunoreactions and chemiluminescence signals recorded digitally using a C-DiGit blot scanner (LI-COR). Band intensities were measured using Image Studio Lite (LI-COR) and data analyzed and graphed with JMP (JMP®, Version 14.0. SAS Institute Inc., Cary, NC, 1989-2019). Blots shown are representative of at least three independent experiments.

### Statistical analysis

Data were analyzed in JMP (JMP®, Version 14.0. SAS Institute Inc., Cary, NC, 1989-2019) or Microsoft Excel. Asterisks represent *p-*value classes as shown. *p*-values were calculated using two-tailed students t-test, other than for quantifications shown in Fig. 1D, which represents one-tailed t-test *p*-values. Graphs were compiled in JMP and typeset using Inkscape (https://inkscape.org/en). Heatmap adapted from GEPIA ^70^.

## Acknowledgments

For plasmids we gratefully acknowledge Drs. R Cohen for Vx3K0-GFP, I Dikic for GFP-OPTN, B van Deurs for GFP-Rab7 and GFP-Rab7^T22N^ and E Holzbaur for GFP-TAX1BP1. We thank the technical staff of the Electron Microscopy Core Facility at Yale School of Medicine and the Johns Hopkins School of Medicine Microscopy Facility for excellent assistance for electron microscopy. This work was supported by Startup funds No. 1606500086 of the W. Harry Feinstone Department of Molecular Microbiology & Immunology, Johns Hopkins Bloomberg School of Public Health (AH-B) and a Johns Hopkins Catalyst Award (AH-B). TSW was supported by the Biochemistry, Cellular, and Molecular Biology graduate program of the Johns Hopkins University School of Medicine, and by the National Institutes of Health (NIH) through Grant No. T32AI007417 (to the Department of Molecular Microbiology & Immunology, Bloomberg School of Public Health).

## Author Contributions

Conceptualization, N.R.B. and A.H.-B.; Methodology, T.S.W., I.C., N.R.B. and A.H.-B.; Formal Analysis, T.S.W., I.C. N.R.B. and A.H.-B.; Investigation, T.S.W., I.C., N.R.B. and A.H.-B.; Data Curation, N.R.B. and A.H.-B.; Writing – Original Draft, N.R.B. and A.H.-B.; Writing – Review & Editing, T.S.W. and I.C.; Visualization, T.S.W., I.C., N.R.B. and A.H.-B.; Supervision, N.R.B. and A.H.-B.; Funding Acquisition, A.H.-B.

