## Supplementary Information for "XIAP-mediated targeting of endolysosomes to stressed mitochondria occurs in a switch-like, global manner and results in autophagy-independent, sub-organelle level mitochondrial degradation"

This PDF file includes:

Supplementary figures 1 to 2

Legends for supplementary movies 1 to 4

Other supplementary materials for this manuscript include the following:

Supplementary movies 1 to 4

### Supplementary Figures

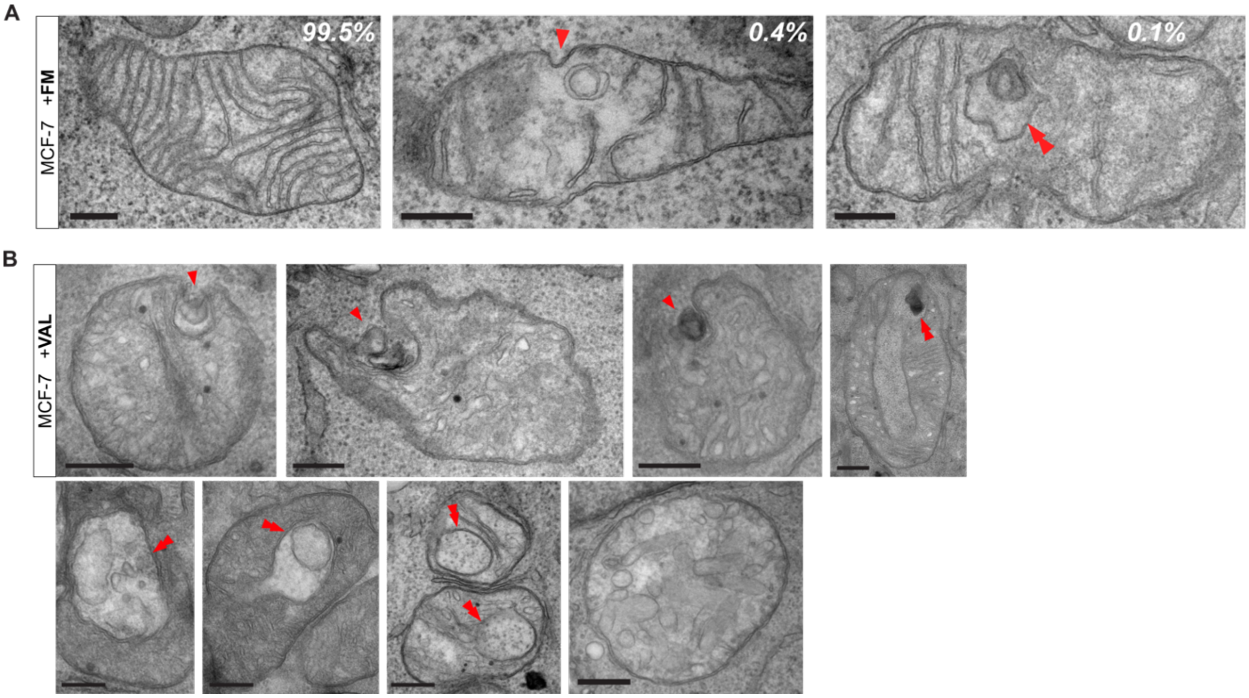


**Supplementary Fig. 1: Ultrastructural analysis of endolysosomal interactions with, and processing of, depolarized mitochondria.**

(A) Transmission electron micrographs of MCF-7 cells kept in control media (full medium, FM). In 99.5% of control cells, mitochondria displayed their typical ultrastructure (*left panel*). 0.4% of control cells presented with endocytic cup-like invaginations in the OMM (arrow, *middle panel*). 0.1% of control cells contained vesicular structures inside inner mitochondrial compartments, surrounded by a double-membrane (arrowheads, *right panel*). Scale bars, 200 nm.

(B) Transmission electron micrographs of MCF-7 cells treated with valinomycin (VAL) for 3 h. Mitochondria oftentimes presented with endocytic cup-like invaginations of the OMM and

OMM-associated IMM, with electron-dense, membrane-bound vesicles (arrowhead) located within these cups. In addition, mitochondria were detected that contained internalized, and double membrane-surrounded (arrowheads), vesicles of various shapes and electron-densities, together with ribosome-containing cytoplasm. Mitochondrial cristae of VAL-treated cells were ultrastructurally altered or absent. Scale bars, 200 nm.

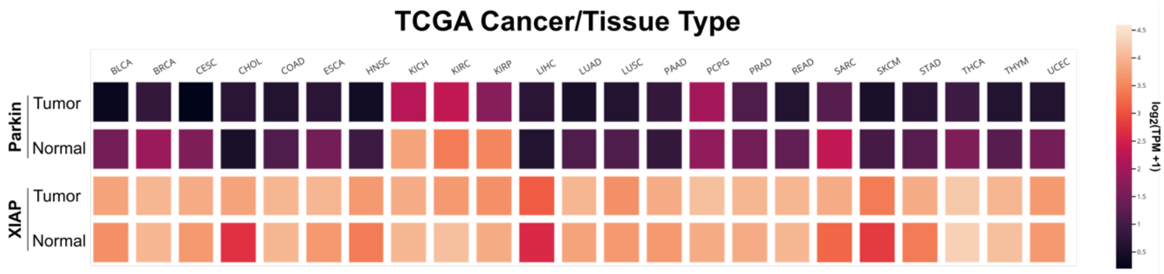


**Supplementary Fig. 2: Heatmap of Parkin and XIAP expression levels.**

Heatmap of Parkin and XIAP expression levels in indicated TCGA cancer and matched normal tissue datasets. Generated using Gepia2 (http://gepia2.cancer-pku.cn/#analysis); expression levels are expressed in $\log_2$(TPM+1).

**Supplementary Movies**

**Supplementary Movie 1: Global, switch-like targeting of Rab5+ endosomes to the mitochondrial compartment following valinomycin-induced depolarization. Related to Fig. 2a.**

Representative movie of MCF-7 cell co-expressing GFP-OMM and RFP-Rab5, imaged *live* every 60 s, starting after 3 h of VAL treatment. Scale bars, 10 μm.

**Supplementary Movie 2: Concurrent XIAP translocation to depolarized mitochondria and largescale mitochondrial accumulation of Rab5. Related to Fig. 2g.**

Representative movie of MCF-7 cells co-expressing GFP-XIAP and RFP-Rab5, *live* imaged every 1 min over 40 min, after VAL administration.

**Supplementary Movie 3: XIAP translocation to depolarized mitochondria and mitochondrial K63-linked polyubiquitylation spatio-temporally coincide. Related to Fig. 2j.**

Representative movie of MCF-7 cells co-expressing RFP-XIAP and K63-linked polyubiquitin chain sensor Vx3K0-GFP, *live* imaged every 1 min over 50+ min, after VAL administration.

**Supplementary Movie 4: Only a small fraction of XIAP-targeted depolarized mitochondria are in proximity with autophagosomes. Related to Fig. 5a.**

Representative movie of MCF-7 cells co-expressing GFP-XIAP (green) and RFP-LC3 (red), *live* imaged every 5 s for a total of 2 min, starting at 3 h of treatment with VAL.
